# Complete mitochondrial genomes of Arizona West Nile virus vectors, *Culex quinquefasciatus* and *Culex tarsalis*

**DOI:** 10.64898/2026.08.14.744795

**Authors:** Zachary A. Barrand, Chase L. Ridenour, Daryn E. Erickson, Alexis N. Rivas, Brooke K. Schmidt, James Will, Steven J. Young, Nicole Busser, John Townsend, Daniel Enriquez, Danyael Murphy, Shukmei Wong, Jonathan Keats, Sophia T. Carvalho, Geoffrey M. Attardo, Christopher M. Barker, Crystal M. Hepp

## Abstract

Here we report a newly developed method utilizing long-range PCR and long-read Pacific Biosciences HiFi sequencing that successfully obtained two full-length and annotated mitochondrial genomes from *Culex quinquefasciatus* Say, 1823 and *Culex tarsalis* Coquillett, 1896, both from Maricopa County, Arizona, USA. Given the substantial burden of West Nile virus in Maricopa County over the past decade, and that these vectors are primarily responsible for spillover to human populations in the county, it is critical to better understand their distribution over time and space. This study begins to approach this need by contributing a novel approach that has resulted in the first West Nile virus vector mitochondrial genomes from Arizona. Our circular *Cx. quinquefasciatus* mitogenome is 15,587 bp in length, making it the first USA-based mitogenome sequenced through the AT-rich control region. The *Cx. tarsalis* mitochondrial genome is 16,416 bp long, longer than recently published California-based CTarK1 and Texas-based PQ585801 mitogenomes. The increased length of the *Cx. tarsalis* mitogenome is a result of a 905 bp insertion in the AT-rich control region, not present in the species’ publicly available mitogenomes. A maximum likelihood-based phylogenetic reconstruction supports the species designation of these newly-sequenced mitogenomes. The newly developed methodology offers a unique approach to study medically-important vector species around the globe, providing a solution to study populations through pooled vector pathogen surveillance programs.

## Introduction

Mosquitoes have been recognized as the most medically-relevant arthropod taxon, responsible for transmission of pathogens including malaria parasites and West Nile (WNV), dengue, yellow fever, chikungunya, and Zika viruses. Since WNV was introduced to the United States in 1999, it has become the most prevalent arbovirus and has become endemic to many regions throughout the country, persisting and circulating between resident bird reservoirs and *Culex* spp. mosquitoes. Specifically, WNV is primarily vectored by the *Culex pipiens* species complex and *Culex tarsalis* Coquillett, 1896 (Fig 1) in the United States (1–4). Due to their medical significance, understanding the distribution and gene flow within these mosquito populations is critical for characterizing the transmission and associated human infection risk of pathogens they transmit.

**Fig 1.**
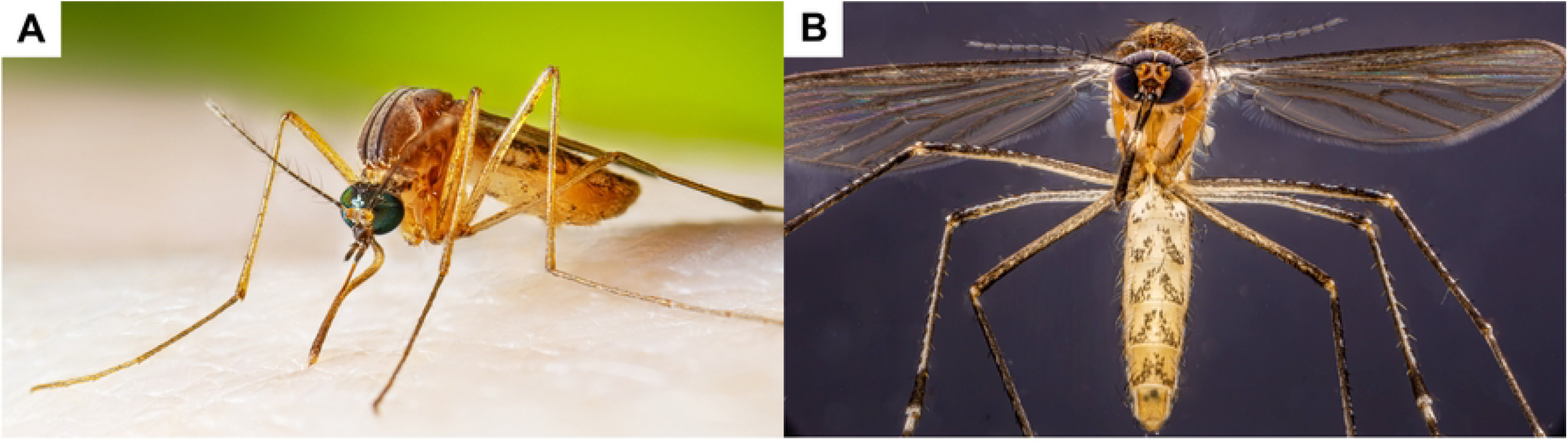
Species reference images. A) Adult female Cx. quinquefasciatus, Lauren Bishop. Publicly-available through the CDC Public Health Image Library, ID# 26024; provided by the CDC and Amy E. Lockwood, MS. B) Adult female Cx. tarsalis, Geoffrey M. Attardo, PhD. Provided by Dr. Attardo from the Department of Entomology and Nematology at UC Davis.

*The Culex pipiens* species complex has been studied extensively throughout the globe, due to their medical significance. Various methods have been used to study these mosquitoes. Typical methods include microsatellites, small genomic or mitochondrial gene segments, or whole genomes using Sanger or Illumina sequencing, respectively (5–7). These methods are usually applied to single mosquitoes. Pooled samples offer a unique way to study these populations as vector control agencies utilize pooled methodology when performing routine pathogen surveillance, but applying widely used methodologies to pooled samples limits data resolution (8,9). For instance, whole-genome data would lead to ambiguities and incorrect genome assemblies and microsatellites, lacking the ability to achieve individual-level allele frequencies from pools (7,9).

To address the need to achieve high-resolution data from pooled samples, we employed a long-range PCR methodology in conjunction with Pacific Biosciences (PacBio) HiFi sequencing to obtain full-length mitochondrial genomes (mitogenome) from *Culex* species mosquitoes. Applying these methods, we obtained the first full-length, publicly available USA-based mitogenome for *Culex quinquefasciatus* Say, 1823 and a third full-length *Cx. tarsalis* mitogenome. Both, from Arizona, USA, were obtained through our novel PCR-based and PacBio HiFi long read sequencing approach.

## Materials and methods

Mosquitoes were collected as part of routine surveillance carried out by the Vector Control Division of Maricopa County Environmental Services, using carbon-dioxide-baited traps (10,11). Individual homogenized *Cx. quinquefasciatus* and *Cx. tarsalis* females were selected for this study. DNA and RNA was co-extracted using the Zymo Research Quick-DNA/RNA Pathogen Miniprep kit. The extracted DNA from the selected *Cx. quinquefasciatus* and *Cx. tarsalis* were deposited at the Museum of Southwestern Biology Division of Genomic Resources at the University of New Mexico (https://msb.unm.edu/divisions/genomics/index.html) under the voucher numbers MSB:DGR:3552 and MSB:DGR:3551 respectively.

Long-range PCR (LRPCR) amplification of whole mitogenomes was performed on extracted DNA using previously published invertebrate primers to amplify *Cx. quinquefasciatus* and *Cx. tarsalis* mitogenomes (S1 Table) (12,13). Amplifications were performed using Promega GoTaq Long PCR Master Mix on a Bio-Rad T100 thermal cycler, with an initial denaturation at 94°C for 2 minutes, 35 cycles of 94°C for 15 seconds, 65°C for 20 seconds, and 65°C for 15 minutes, and a final 10-minute extension at 72°C. Amplification product was held at 4°C until further analyses to avoid freeze-thaw of amplified DNA. Each 25µL PCR reaction contained primers with attached PacBio M13 tails, at concentrations of 300 nM. The product was observed on a 0.5% agarose gel with New England Biolabs Quick-Load 1 kb Extend DNA Ladder, ensuring a target size of ∼15kb (S1 Fig).

Long-range amplifications were cleaned with AMPure PB PCR purification beads (PacBio) at 0.4X and eluted in PacBio Elution Buffer. Samples were barcoded and further prepared using a modified version of the PacBio protocol PN 101-921-300. Each barcoding PCR reaction consisted of 12.5µL of Promega GoTaq Long PCR Master Mix 2X, 2.5µL of Barcoded M13 primer pair (unique to each sample), 6.5µL of cleaned gene-specific PCR product, and 3.5µL molecular biology-grade water for a final volume of 25µL. Barcode PCR conditions were: initial denaturation of 94°C for 2 minutes, 10 cycles of denaturation at 94°C for 15 seconds, annealing at 60°C for 20 seconds, and extension at 65°C for 15 minutes, and a final extension at 72°C for 10 minutes. Following barcoding, samples were cleaned as described above and quantified using a high-sensitivity Invitrogen Quant-iT dsDNA Assay Kit on a BioTek plate reader. Resulting fluorescence values from kit standards were used to construct a standard curve to calculate nanomolar concentration of cleaned samples, which were pooled equimolar into a final library and quantified using a high-sensitivity Invitrogen Quant-iT dsDNA Assay Kit on a Qubit 2.0 instrument to ensure adequate concentration for downstream library preparation using the PacBio protocol PN 101-921-300. Genomic DNA TapeStation Analysis was performed on an Agilent Technologies TapeStation 4200 system to ensure only desired DNA fragments were present. Quantified libraries were sent to the TGen Collaborative Sequencing Center for PacBio Revio sequencing.

Demultiplexed sequence reads underwent primer trimming in Lima v2.9.0 followed by BAM to fastq conversion using the PacBio bam2fastq tool. Due to the presence of non-specific amplicons, reads were classified with Kraken2 v2.1.3 using the core nucleotide database (14). The reads classified to *Culex* spp. were extracted using the kraken extract tool, removing unwanted reads. Minimap2 v2.26 was used for reference-based alignment of the extracted classified reads, with MN389462 as the *Cx. quinquefasciatus* reference and the CTarK1 mitogenome as the *Cx. tarsalis* reference (15–17). Consensus mitogenomes were obtained in iVar v1.4.3, where sites were called if the majority allele frequency was at least 80% (18).

Following assembly, the Culex tarsalis mitogenome contained a stretch of 1,800 ambiguous bases due to back-to-back misalignment of a 905 bp repetitive AT-rich insertion. To achieve the correct alignment of the insertion, we first generated our own *Cx. tarsalis* reference mitogenome. To do this, we aligned all of the reads over 15kb using MAFFT v7.525. The resulting alignment was imported into Geneious Prime v2025.2 and a consensus was generated at a 90% threshold, ignoring gaps. The orientation of this consensus mitogenome was then corrected using the CTarK1 reference and the Kraken2 classified reads were then mapped to the newly made reference mitogenome using Minimap2 v2.26. The final consensus mitogenome was obtained in iVar v1.4.3 at an 80% threshold. Annotation and circularization of both mitogenomes was carried out using a set of reference genomes (EU352212, HQ724617, KT851543, KT851544, MF509895, MN389459, MN389462, AY072044, HQ724614, MF040164, MF381652, MF381736, KX753344, MN389456, OL351548) in Geneious Prime v2024.0.2 (Fig 2).

**Fig 2.**
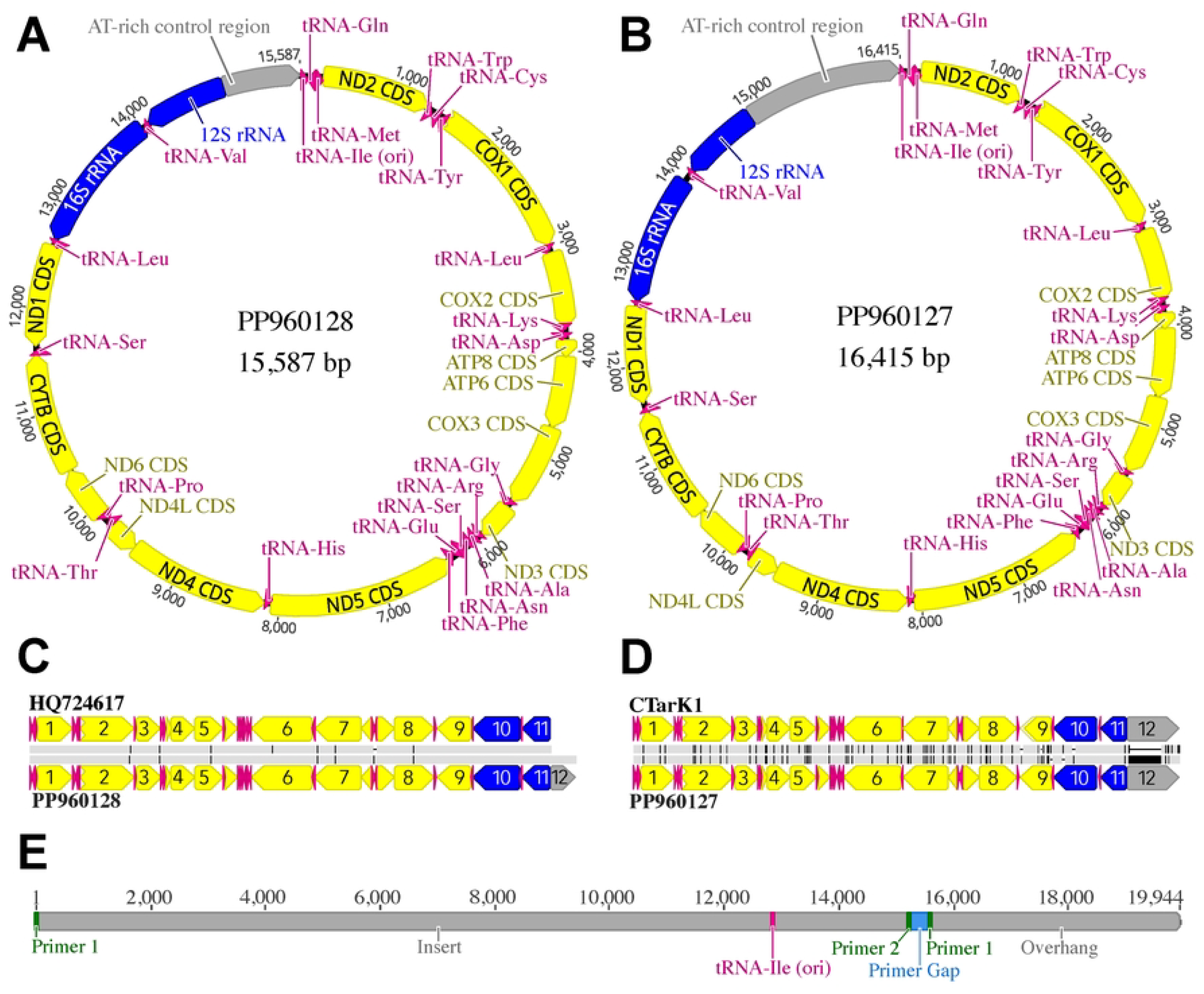
Genome feature map of Cx. quinquefasciatus (A) and Cx. tarsalis (B) mitochondrial genomes through reference-based annotation in Geneious Prime v2024.0.2. Each has 13 protein-coding genes (yellow), 22 transfer RNA genes (pink), two rRNA genes (blue), and a non-coding AT-rich control region (gray). The origin (ori) is noted at the tRNA-Ile (pink). (C) Linear alignment of USA-based Cx. quinquefasciatus mitogenome reference HQ724617 and newly obtained Cx. quinquefasciatus mitogenome PP960128. (D) Linear alignment of USA-based Cx. tarsalis mitogenome reference CTarK1 and newly obtained Cx. tarsalis mitogenome PP960127. Linear mitogenomes start at the ori (tRNA-Ile) with the numbered genes in the following order: (1) ND2, (2) COX1, (3) COX2, (4) ATP6, (5) COX3, (6) ND5, (7) ND4, (8) CYTB, (9) ND1, (10) 16S rRNA, (11) 12S rRNA, and (12) AT-rich control region. (E) Annotated raw PacBio HiFi reverse read for Cx. quinquefasciatus, starting at the reverse primer binding site (Primer 1) within the 16S rRNA gene at positions 12,891 – 12,858 for the resulting Cx. quinquefasciatus mitogenome (A). The long read indicates a full-length mitogenome was obtained via amplification around the circular target through the forward primer binding site (Primer 2) and back to the reverse primer binding site (Primer 1) (E).

To confirm species placement of our assembled mitogenome, we performed phylogenetic comparisons using one representative mitogenome per Culex species from GenBank, with *Aedes aegypti* and *Aedes albopictus* as outgroups. Additional mitogenomes were pulled from GenBank for the *Cx. pipiens* complex and *Cx. tarsalis* to account for geographic differences, including the CTarK1 mitogenome provided by Bradley Main and Christopher Barker (15,16). Additionally, *Cx. amazonensis* mitogenome was provided by Alexandre Freitas da Silva and included in phylogenetic analysis (19). The full-length mitogenome dataset, from origin (ori) through the AT-rich control region, was aligned using MAFFT v7.525. The resulting alignment had a pairwise percent identity (S2 Fig) and a pairwise number of differences (S3 Fig) calculated in Geneious Prime v2025.2, followed by model selection in IQTree v2.3.6 (20) with ModelFinder. The GTR+F+I+R3 evolutionary model had the best fit to the full-length alignment given the Bayesian Information Criterion. Phylogenetic reconstruction using maximum likelihood and ultrafast bootstrapping was also performed in IQTree with 10,000 bootstrap replicates. Phylogenetic visualization was carried out using FigTree v1.4.4.

## Results and discussion

Two mosquitoes were selected for our study. One *Cx. quinquefasciatus* and one *Cx. tarsalis* from Queen Creek, Arizona (Latitude: 33.255, Longitude: -111.607) and Surprise, Arizona (Latitude: 33.643, Longitude: -112.456), respectively (Fig 1). Our LRPCR methodology resulted in amplification of our target 15kb amplicon as well as additional fragments of varying length, including several small fragments <3kb and smearing above 15kb (S1 Fig). The small amplicons were likely generated from either (1) fragmented mitochondrial DNA that was generated during sample and/or nucleic acid handling or (2) off-target/nonspecific amplification where the PCR primers bound to other mtDNA or gDNA. Additionally, smearing was present at the top of the 15kb band, indicative of a variety of larger amplicons migrating through the gel, preventing a clean band from forming (S1 Fig). However, this could also be observed if there was a large amount of product that was not separating properly in the gel due to a high voltage and a fast run time. This was likely not the case, since we used a low percentage agarose at 110V for 1 hour 45 minutes, allowing for appropriate DNA fragment separation. The streaking of our 15kb amplicon band in our agarose gel was likely DNA amplicons of varying length around our 15kb target.

The sequencing run resulted in a variety of read lengths, which is typical of long-read sequencing (S4 Fig) (21). The minimum read lengths observed were 679 bp and 285 bp for *Cx. quinquefasciatus* and *Cx. tarsalis*, respectively (S2 Table). The resulting maximum sequence lengths for *Cx. quinquefasciatus* and *Cx. tarsalis* were 19,944 bp and 19,410 bp respectively, larger than our target size of 15kb (S2 Table). The long reads greater than our target size of 15kb allowed us to achieve full-length mitogenomes. When mapping the reads to the samples, we observed reads that contained our forward and reverse primer sites in the middle of the read, such as the 19,944 bp *Cx. quinquefasciatus* read (Fig 2). We hypothesize these reads were generated from amplification around the circular mitochondrial genome during the initial cycles of LRPCR. However, some short reads mapped to the gap between the primer positions. We hypothesize these short reads spanning this gap were once part of a larger amplicon that then sheared during library preparation.

We have generated full-length mitogenomes of Arizona *Cx. quinquefasciatus* and *Cx. tarsalis* through LRPCR amplification and PacBio Revio sequencing, that are 15,587 bp and 16,416 bp in length at coverage depths of 1,481X and 286X respectively (Fig 2) (S5 Fig). These mitogenomes contain 13 protein-coding genes, 22 transfer RNA genes, two ribosomal RNA genes, and a non-coding AT-rich control region of 730 bp and 1,559 bp in *Cx. quinquefasciatus* and *Cx. tarsalis*, respectively. The structure of our mitogenomes corresponds to the structure previously described for mosquitoes (22,23) (Fig 2). However, these newly obtained mitogenomes are the longest characterized for *Cx. tarsalis*, globally, and *Cx. quinquefasciatus* in the USA to date, due to the presence of an insertion and the inclusion of the AT-rich control region respectively (Fig 2). During mitogenome assembly, a 905-bp insertion present in the first 80 bp of the AT-rich control region of *Cx. tarsalis* was found in 94.76% (271/286) of the reads over 15kb. Investigating the insertion further, it is a repetitive section of the same ∼287-bp motif. Due to the AT-rich and repetitive nature of the insertion, we faced difficulty aligning the insertion properly using a traditional mapping approach which was applied to the initial mitogenome assembly (PP960127.1) (Fig 3).

**Fig 3.**
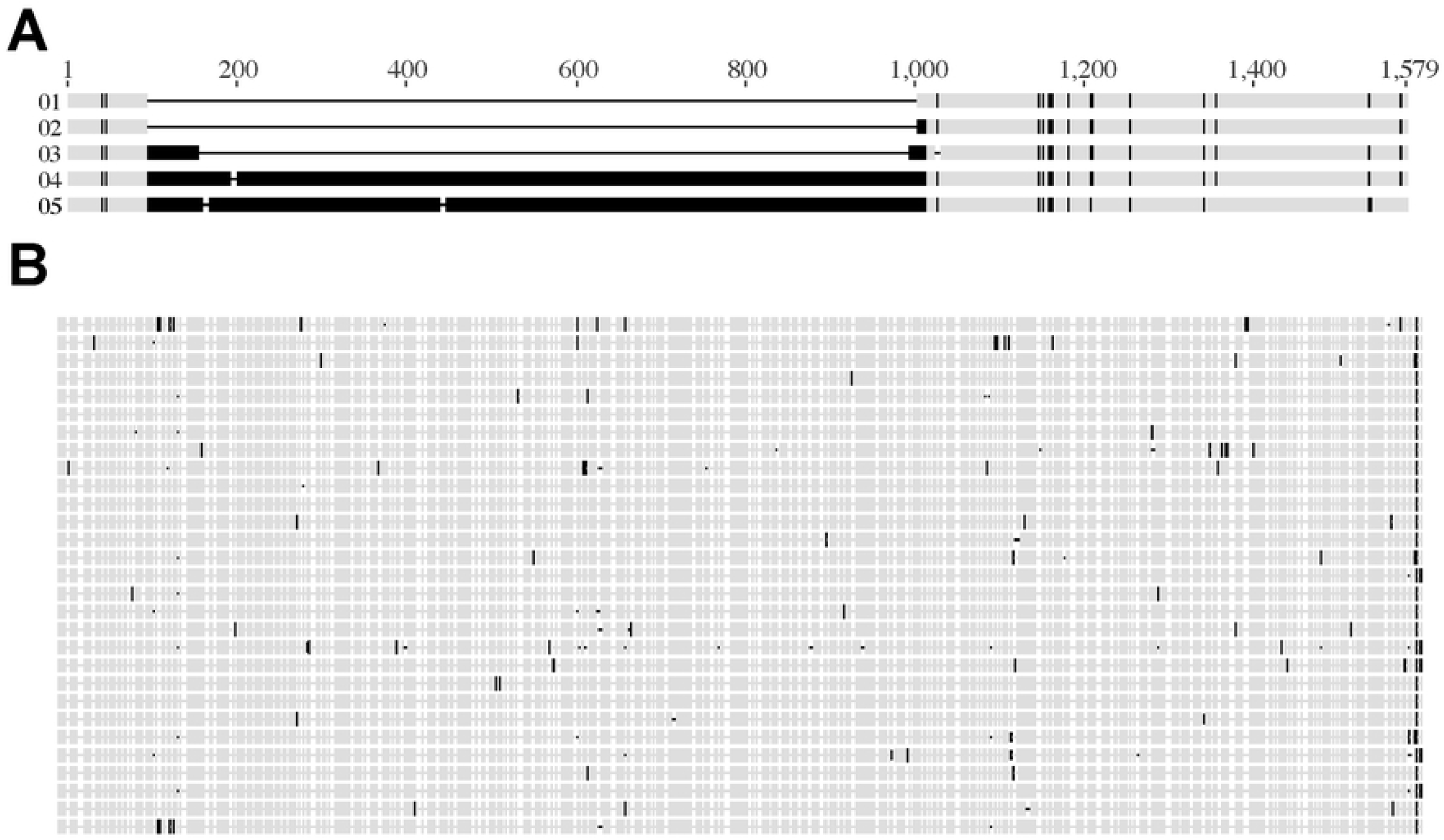
AT-Rich Control Region Alignment. (A) Alignment of five Culex tarsalis AT-Rich Control Regions; CTarK1 (01), PP960127.1 (02), PQ585801 (03), New CTarK1 (04), and PP960127.2 (05). Gray bases indicate similarities in the alignment and Black indicate disagreements in the alignment. CTarK1 and the original PP960127.1 AT-Rich Control Regions do not contain the 905 bp insertion that was discovered in the long-read sequence data. PQ585801 contains more of the insertion than the other two. PP960127.2 contains the full 905 bp insertion in the AT-Rich Control Region. The New CTarK1 mitogenome contains 618 bp of the insertion, 278 Ns are present due to the presence of a 278 bp deletion from PP960127.2. (B) Raw read alignment of the AT-Rich Control Region insert of PP960127.2 from the long-read sequence data. Gray bases indicate similarities in the alignment and Black bases indicate disagreements in the alignment. In the read alignment, there is variability present throughout the region, indicating difficulty aligning the region with traditional mapping approaches.

Comparing our *Cx. tarsalis* mitogenome containing the 905-bp AT-rich control region insertion to other *Cx. tarsalis* mitogenomes, the insertion is not present in the California-based CTarK1 and Texas-based PQ585801 mitogenomes (Fig 3A). However, the Texas-based *Cx. tarsalis* contained more of the bases from the insertion than the CTarK1 mitogenome, which contained none of the bases from the insertion. While we are the first to report this 905-bp mitogenome insertion in the *Cx. tarsalis* AT-rich control region, we do not think it is unique to *Cx. tarsalis* found within Arizona. We hypothesize that the use of Illumina short-read sequencing and/or traditional bioinformatic approaches result in improper assembly of the AT-rich control region. These methods can result in read misalignment and assembly inaccuracies due to the insertion’s AT-rich repetitive nature and length, which is longer than the capabilities of Illumina sequencers. To test this, we obtained the PacBio read data for CTarK1 from Chris Barker at UC Davis and reassembled the mitogenome with our new reference. Following assembly, we generated a new 16,416 bp long CTarK1 mitogenome, including 907 bp of added total length in the AT-rich control region compared to its original iteration (Fig 3A).

Our phylogenetic analysis included a sample of 24 *Culex* spp. mitogenomes, with *Cx. amazonensis* placed in the most basal position as previously observed (19) (Fig 4). The Arizona *Cx. tarsalis* clustered as expected and were most closely related to the *Cx. tarsalis* mitogenome PQ585801 from Harris County, TX (Fig 4). The Arizona *Cx. quinquefasciatus* clustered as expected within the *Cx. pipiens* complex, most closely related to a *Cx. quinquefasciatus* from California, USA (HQ724617) (Fig 4). However, the *Cx. pipiens* complex did not demonstrate strong species clustering in some cases and tended to have lower bootstrap values (Fig 4). We theorize two possible explanations for the unexpected lack of clustering: (1) morphological differences within the *Cx. pipiens* complex are notoriously difficult to distinguish, which could lead to species misidentification (24,25) or (2) hybridization between *Cx. pipiens pipiens* and *Cx. quinquefasciatus*, leading to paternal leakage of mtDNA when there is high genetic diversity (26,27).

**Fig 4.**
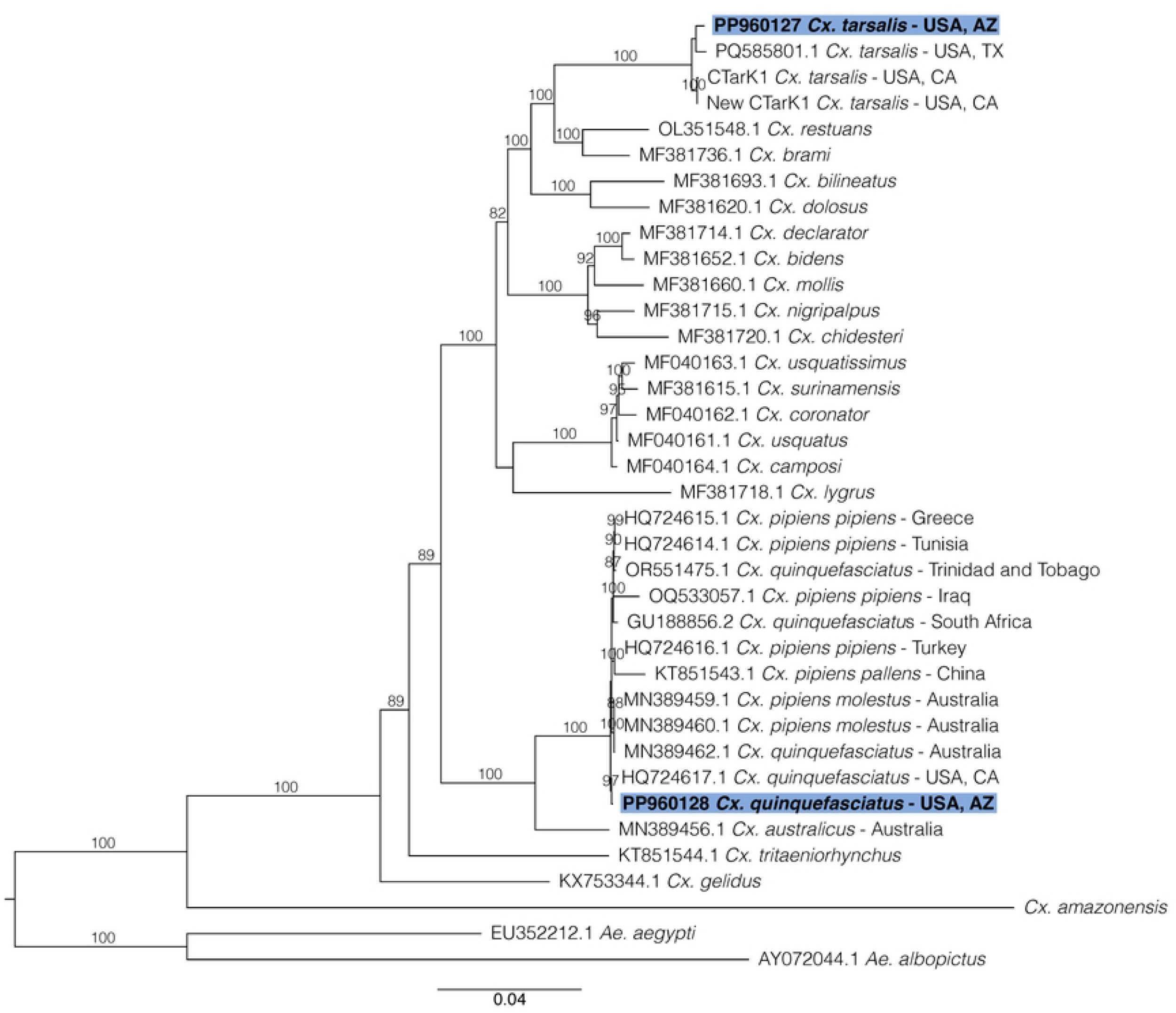
Maximum likelihood phylogeny of full-length mitochondrial genomes (mitogenome) from 24 Culex spp. mosquitoes. Aedes aegypti and Ae. albopictus are used as the outgroup. The mitogenomes sequenced as part of this study are bolded with a blue background. Collection countries are listed for the Culex pipiens spp. complex to highlight geographic rather than species clustering. The scale bar gives a substitutions per site indication. Bootstrap values are provided for branches with a bootstrap value greater than or equal to 80.

## Conclusion

Here we have introduced a long-range PCR approach coupled with Pacific Biosciences HiFi sequencing to obtain full-length mitochondrial genomes from *Cx. quinquefasciatus* and *Cx. tarsalis*. The two new *Cx. quinquefasciatus* and *Cx. tarsalis* mitogenomes presented here as a result of our approach are the longest to date for both species in the United States. Our approach is significant, offering a solution to achieve high-resolution data, expanding the potential of population-level studies through routine vector control surveillance programs utilizing pooled samples. Monitoring the interactions within mosquito populations such as *Cx. quinquefasciatus* and *Cx. tarsalis* can provide deeper insight into the maintenance and circulation of the pathogens they transmit, such as WNV. The new approach outlined here has the potential to aid in understanding medically-significant species around the globe, offering further insight into vector and pathogen movement and in turn informing vector control mitigation strategies.

## Supporting Information

**S1 Table. Previously published long-range PCR primers for amplification of completed arthropod mitochondria**. Doi: 10.1006/mpev.2001.0940

**S1 Fig. 0.5% agarose gel image of mitogenome Long-Range PCR product for Culex quinquefasciatus**

**(2) and Culex tarsalis (3)**. Three PCR No Template Controls were cycled alongside the samples (4,5,6). The New England Biolabs Quick-Load 1 kb Extend DNA Ladder was used to visualize the high molecular weight amplification product (1). The ladder is annotated in kilobases.

**S2 Fig. Pairwise percent identity matrix of aligned Culex (n=34) and Aedes (n=2) mitochondrial genomes contained in the maximum likelihood phylogeny**.

**S3 Fig. Pairwise number of differences matrix of aligned Culex (n=34) and Aedes (n=2) mitochondrial genomes contained in the maximum likelihood phylogeny**.

**S4 Fig. *PacBio HiFi read length histogram of (A) Cx. quinquefasciatus and (B) Cx. tarsalis***.

**S2 Table. Minimum and Maximum lengths for raw PacBio HiFi reads**.

**S5 Fig. Genome coverage plot of (A) Cx. quinquefasciatus and (B) Cx. tarsalis mitogenomes with mitogenome position (x-axis) versus the log transformed read count (y-axis)**. The first position of the plot is associated with position 13,221 of the Cx. quinquefasciatus reference MN389462 and position 13,211 of the previously published Cx. tarsalis genome.

**S1 File. Mosquito sample metadata for the newly obtained mitochondrial genomes. S2 File. Mosquito mitogenome nucleotide alignment**.

